# SOX2 and SOX9 as Transcriptional Regulators of human Galectin-3 in SW1353 Cells: Potential Implications for Osteoarthritis

**DOI:** 10.1101/2025.06.03.657610

**Authors:** Blanca Alba, Herbert Kaltner, Stefan Toegel, Sebastian Schmidt

## Abstract

Galectin-3 (Gal-3), a member of the β-galactoside-binding protein family, is critically involved in inflammation, extracellular matrix remodelling, and cartilage degeneration in osteoarthritis (OA). This study aims to elucidate the regulation of the human galectin-3 gene (*LGALS3*) promoter in SW1353 cells and its control by SOX transcription factors, known to be dysregulated during OA pathogenesis. We sought to identify key sequence elements in the *LGALS3* promoter responsible for its transcriptional activity and the transcription factors (TFs) responsible for its regulation. Using luciferase reporter assays, we examined deletion variants of the 5’ region (-2638 bp to +52 bp) and assessed their activation potential. We also identified potential transcription factor binding sites (TFBS) through *in silico* analyses and confirmed SOX9 binding in the -93/+49 region by chromatin immunoprecipitation using HaloCHIP^TM^. Functional assays revealed that the proximal promoter region (-97 to +52 bp) is critical for reporter gene expression in SW1353 cells. This study demonstrates that the presence of *SOX2* and *SOX9* leads to a dose-dependent decrease in *LGALS3* promoter activity in SW1353 cells.

We show that SOX9 can bind the promoter, highlighting the importance of SOX TF interactions in regulating *LGALS3* expression and their potential role in chondrocytes.

## 1. Introduction

Galectins are an evolutionarily conserved family of glycan-binding proteins characterized by a conserved β-galactoside-binding carbohydrate recognition domain (CRD) consisting of a seven-amino acid sequence signature [1]. The 3D conformation of CRDs is “a β-sandwich described as a jelly-roll motif” [1]. Galectins are abundant across diverse species, present in vertebrates, protochordates, invertebrates, mushrooms, and viruses [2,3].

Galectins can be classified according to the arrangement of their CRDs into three subgroups: prototype, tandem-repeat type, and chimera type. The chimera type galectin, represented solely by galectin-3 (gene name: *LGALS3*), possesses a unique structure combining a C-terminal CRD with an N-terminal domain rich in glycine-proline repeats, which facilitates oligomerization [1]. Galectin-3 levels in patients’ sera are characteristic of both osteoarthritis (OA) and early rheumatoid arthritis, underscoring the relevance of understanding the regulatory mechanisms of *LGALS3* in both conditions [4–6]. In OA, a degenerative joint disease characterised by chronic, low-grade inflammation [7–9], galectin-3 presence in articular chondrocytes correlated positively with the severity of cartilage destruction [10]. Furthermore, extracellular galectin-3 bound to the surface of OA chondrocytes and induced the expression of pro-inflammatory cytokines and matrix metalloproteases by activating the NF-kB pathway [10]. This cascade eventually resulted in the degradation of extracellular matrix in a 3D organoid model of OA chondrocytes [11]. However, the exact mechanisms that result in the observed upregulation of galectin-3 in human OA chondrocytes are still unexplored, underscoring the need for studies that deepen our understanding of its molecular regulation in this disease condition.

*LGALS3* expression can be regulated by transcriptional and epigenetic mechanisms [12,13]. Among these regulatory layers, transcriptional control is particularly relevant for understanding how *LGALS3* expression is altered in OA, since extensive transcriptomic changes in chondrocytes occurred in progressing OA including significant alterations in transcription factor expression [14]. Crucial transcription factors in chondrocytes belong to the SOX (SRY-related HMG-box) protein family, whose members have gained particular attention as master regulators of differentiation. There are 20 SOX transcription factors conserved in mammals. The consensus sequence recognized by most SOX transcription factors is 5’- (A/T)(A/T)CAA(A/T)G -3’ [15]. SOX proteins can bend the DNA upon binding to allow for chromatin modelling. They are crucial in cell fate determination, differentiation, and development [15]. SOX9, widely recognised as ‘the master regulator of cartilage development’, is critical in cartilage homeostasis, activating the expression of extracellular matrix (ECM) components such as collagen type II (COL2A1) and aggrecan (ACAN) [16]. SOX9, together with SOX5 and SOX6, build the SOX trio [16]. These three SOX proteins are downregulated in OA chondrocytes [17], whereas another member, SOX4, is upregulated [18]. Also, SOX9 induces the expression of other *SOX*, i.e., *SOX5* and *SOX6* [16].

Another member of the galectin family, galectin-1, is reported to activate the β-catenin-SOX9 signalling cascade, thereby promoting tumour progression in colorectal cancer [19] and showing a potential linkage of SOX proteins and galectins. Despite the well-established role of SOX transcription factors in cartilage homeostasis and OA progression, their direct involvement in regulating galectins remains unexplored. Here, we hypothesise that SOX factors impact the expression of *LGALS3* in human chondrocytes.

Therefore, this study aims to identify the key regulatory elements of the *LGALS3* promoter and SOX-binding motifs and assess the impact of members of the SOX family on gene expression in SW1353 cells, a human chondrosarcoma-derived cell line commonly used as a chondrocyte model [20–22]. By analysing *LGALS3* promoter deletions, we identified the minimal promoter region with maximal activity. Overexpression experiments revealed that *SOX2* and *SOX9* dose-dependently reduced promoter activity to 10 %, whereas *SOX4*, *SOX5,* and *SOX6* had no significant influence. This repression with *SOX2* and *SOX9* persisted even with the shortest promoter construct (-97/+52). This study is the first to demonstrate that SOX transcription factors directly regulate the *LGALS3* promoter, establishing a novel link between the SOX family and galectin-3 expression in chondrocytes.

## 2. Methods

### 2.1. Chondrocyte cell culture

SW1353 human chondrosarcoma cells were cultured at 37 °C in a humidified atmosphere with 5 % CO2 in Dulbecco’s Modified Eagles Medium (DMEM), containing 10 % FCS and 1 % penicillin/streptomycin/L-glutamine. Primary OA chondrocytes were isolated from human articular knee joint cartilage obtained from OA patients during total knee replacement surgery with written informed consent and following the terms of the ethics committee of the Medical University of Vienna (EK-No. 1822/2017 and 1555/2019). Primary chondrocytes were cultured under standard conditions [23] and used without subculturing to preserve the chondrocyte phenotype.

### 2.2. Plasmid extraction

Plasmid extraction was performed with GeneJET Plasmid-MiniPrep-Kit (Thermo Fisher Scientific, Dreieich, Germany) according to the manufacturer’s instructions. The purified plasmid was eluted in 50 µL of water to be able to use the plasmid for sequencing and further downstream applications. The plasmids were obtained from E. coli TOP10 chemically competent cells (Thermo Fisher Scientific). The plasmids used are listed in Supplementary Table S1.

### 2.3. RNA isolation and quantitative PCR (qPCR) analysis

Total RNA was isolated from primary OA chondrocytes and SW1353 cells using the Invitrogen™ RNA Mini Kit (Thermo Fisher Scientific) following the manufacturer’s instructions, adding a DNase digestion step to avoid genomic DNA contamination. For RNA extraction from transfected cells, the SW1353 cells were cultured in a 6-well plate and harvested 48 hours post-transfection. For cDNA synthesis, 2 µg of the isolated RNA was reverse transcribed using the GoScript™ Reverse Transcription System (Promega, Walldorf, Germany). RNA concentration and purity were assessed using a Nanophotometer IMPLEN® P300 (IMPLEN, Munich, Germany), with A260/A280 ratios between 1.98 and 2.07 indicating high purity. RNA integrity was evaluated by agarose gel electrophoresis, confirming the presence of intact rRNA bands. For qPCR, we used 10 μL of LUNA® Universal qPCR Master Mix (NEB, Frankfurt am Main, Germany) in a final reaction volume of 20 μL. Reactions were carried out in technical triplicates on the CFX Duet Real-Time PCR System (Bio-Rad, Munich, Germany), and data were processed using the Bio-Rad CFX Maestro software. Real-time PCR primers were designed manually, checking for primer-dimer formation and secondary structures using the OligoEvaluator™ tool (https://www.oligoevaluator.com/LoginServlet). Primer sequences are presented in Supplementary Table S2. Primers were designed to span exon-exon junctions, to minimize genomic DNA amplification when possible. Amplification efficiencies were determined for each primer pair from standard curves using serial dilutions of cDNA, and only primer sets with efficiencies between 106-112% and R² values close to 1 (Supplementary Table S3) were included in the analysis. Melt curve analysis confirmed a single peak, meaning specific amplification products. Relative gene expressions of GAPDH, LGALS3, SOX2, SOX4, SOX5, SOX6, and SOX9 were calculated using the Pfaffl method [24], using GAPDH as the reference gene.

### 2.4. Construction of human LGALS3 promoter reporter gene vectors

The promoter sequence was amplified from the genomic DNA of SW1353 cells. We used the Wizard® Genomic DNA purification kit (Promega) to extract the genomic DNA. We amplified the first fragment of the promoter (hGal3p-I) by using Expand™ Long Template PCR-System (Roche, Mannheim, Germany). The second fragment (hGal3p-II) was amplified by GC-RICH PCR-System (Roche). Both fragments were first subcloned into pGEM-T Easy (Promega), then digested with NheI and EcoRI (hGal3p-I) or EcoRI and HindIII (hGal3p-II). The resulting fragments were then ligated in a three-fragment ligation into pGL4.20 that was cut using NheI and HindIII and dephosphorylated using the rAPid Alkaline Phosphatase (Roche) to create the pGL4-hGal3p(-2636/+52) reporter vector. Sequence deletions of the LGALS3 promoter were cloned accordingly using GC-RICH PCR-System (Roche) or Phusion® High-Fidelity DNA Polymerase (NEB). Primer sequences used are in Supplementary Table S2. The sequences of the constructs were all verified by commercial sequencing.

### 2.5. Construction of transcription factor expression vectors

The coding sequences of the human transcription factors SOX2, SOX4, SOX5, and SOX6 were amplified by PCR from cDNA of HEK293 cells. The sequence of SOX9 was amplified from a commercial plasmid, FUW-tetO-SOX9, which was a gift from Lorenz Studer (Addgene, Watertown, US; plasmid #141404; RRID:Addgene 141404) [25] using the primer sets listed in Supplementary Table S2. The amplicons were digested with the appropriate restriction enzymes and directly ligated into the expression vector pcDNATM3.1(+) (Thermo Fisher Scientific). Plasmids were commercially sequenced to verify the inserted sequences.

### 2.6. Cell transfection

For transfection, SW1353 cells were grown until 70–80 % confluency. A volume of 0.5 mL of a 2 x 105 cells/mL cell suspension was seeded into each well of a 24-well cell culture plate. Alternatively, 2 mL of this cell suspension were seeded in the experiments using 6-well plates. After incubation for 24 h, the cells in 24-well plates were transfected with 500 ng of DNA per well using the PEI STAR™ (Tocris Bioscience™, Bristol, UK) transfection reagent in a ratio of 2 μL of PEI STAR™ (1mg/mL) per 1 μg of DNA according to the manufacturer’s protocol. Similarly, the cells in 6-well plates were transfected with 2 µg of DNA following the same PEI STAR™ (Tocris Bioscience™) ratio.

The transfection mixtures for luciferase reporter assays contain 50 ng of pGL4-hGal-3 promoter plasmid, 10 ng pGL4.74 (Renilla luciferase) for normalization and for the overexpression experiments, increasing amounts (ranging from 0.1 to 100 ng) of the transcription factor-expressing pcDNA3.1(+) plasmid. Each mixture was supplemented with an empty pGEM®-T Easy vector (Promega) to reach 500 ng DNA per 24-well in all transfection tubes, ensuring equal DNA amounts. The culture plate was gently agitated, and the cells were grown for 24 h before further processing, adding 500 µL of media after 2 hours to avoid toxicity from the transfection reagent. For the 48h overexpression followed by qPCR analysis, we used 3 µg of SOX-encoding pcDNA3.1(+) within a 6-well.

### 2.7. Luciferase reporter gene assays

Luciferase measurements were carried out using 20 μl of the SW1353 cell lysate and following the instructions of the Dual-Luciferase® Reporter Assay System (Promega). The readout of the signals was obtained using the plate reader Infinite 200 (Tecan, Crailsheim, Germany) with an integration time of 5 s for firefly and Renilla luciferase. The firefly/Renilla signal ratio was calculated to normalize the signal according to the transfection efficiency.

### 2.8. Sequence alignment (mVISTA and Jalview software)

For the sequence alignments, we searched for the LGALS3 orthologues in a selection of species representing different taxonomic groups: Rhesus monkey (Macaca mulatta, Gene ID: 697290), orang-utan (Pongo abelii, Gene ID: 100453870), mouse (Mus musculus, Gene ID: 16854), rat (Rattus norvegicus, Gene ID: 83781), dog (Canis lupus familiaris, Gene ID: 404021), chicken (Gallus gallus, Gene ID: 373917), pig (Sus scrofa, Gene ID: 100038033), horse (Equus ferus caballus, Gene ID: 100050367) and zebrafish (Danio rerio, Gene ID: 557373). Subsequently, all sequences were standardized to a length of 2690bp upstream the translational start ATG and aligned to the 2690bp long human sequence (Homo sapiens, Gene ID: 3958) using the mVISTA software [26,27] (available at https://genome.lbl.gov/vista/mvista/submit.shtml). For a more detailed analysis, we performed a multiple sequence alignment of the -77 to -43 region in Jalview (version 2.11.4.1) using the MAFFT algorithm, and results were viewed using the same software.

### 2.9. Computational analyses

JASPAR software (10th release (2024) https://jaspar.elixir.no) [28], with a percentage threshold indicated in the different analyses, was used to identify potential transcription factor-binding sites. qPCR data were analysed using the ΔΔCt method or the Pfaffl method [24]. The ΔΔCt method was applied for comparing gene expression between the SW1353 cell line and the OA chondrocytes to characterise their SOX2, SOX4, SOX5, SOX6, SOX9, and LGALS3 expression profiles. In contrast, the Pfaffl method was used to calculate the relative gene expression ratio, considering the primer efficiencies. To calculate ΔCt, we normalised the Ct values of the target genes to those of GAPDH. For ΔΔCt values, the ΔCt values of the target genes in primary OA chondrocytes were related to those in the SW1353 cells.

### 2.10. HaloCHIPTM - Chromatin Immunoprecipitation

We used the pcDNATM3.1(+)-SOX9 vector as a template to clone SOX9 into the pHTN vector (Promega), resulting in the plasmid pHTN-SOX9. This vector encodes for a fusion protein which consists of an N-terminal Halo-Tag (33 kDa) and the SOX9 protein. SW1353 cells were transfected within a 6-well plate using 1.5 μg of these vectors and 3 μL of PEI STAR™ transfection reagent. After 24 h at 37 °C and 5 % CO2, cross-linking was carried out using DMEM containing formaldehyde at a final concentration of 1 %. After incubation for 10 min at room temperature, the reaction was quenched by adding 1.25 M glycine (pH 7.0) to a final concentration of 125 mM within the medium, which caused a yellow colour. After washing the cells with PBS, they were scraped and centrifuged at 2,000xg for 5 min at 4°C, and the pellets were frozen for further processing. Subsequent steps followed the manufacturer’s protocol (HaloCHIPTM System, Promega).

### 2.11. Statistical procedures

Statistical comparisons between groups were performed using GraphPad Prism software version 10.0.2 (GraphPad Software, Boston, MA, USA). The non-parametric Kruskal-Wallis test followed by Dunn’s multiple comparisons test was used. Results were statistically significant at *p <0.05, **p <0.01, and ***p <0.001.

## 3. Results

### 3.1. Deletion analysis of the *LGALS3* promoter region

The first aim of this work was to identify critical sequence regions responsible for mediating *LGALS3* gene expression in the SW1353 chondrocyte model. The sequence region we investigated spans between -2638 and +52 relative to the transcription start site (TSS). A series of promoter fragments were inserted into the promoter-less luciferase reporter vector pGL4.20, upstream of the luciferase gene, to generate the different pGL4.20-hGal3p reporter plasmids. The empty promoter-less vector (pGL4.20) served as a reference for calculating the promoter activities.

Figure 1A shows the relative lengths of each promoter deletion construct drawn to scale. Figure 1B shows maximum activities for the sequence stretches (-1442/+52) with 162.9-fold and (- 97/+52) with 135.6-fold activity, and an activity decrease between (-1442/+52) and (-535/+52) to 81.1-fold (Fig. 1B). The promoter activity remains roughly constant between the consecutive deletion constructs from (-535/+52) to (-117/+52). When comparing the full-length promoter (- 2638/+52) with the (-77/+52) deletion variant, the reporter assay activity decreased from 62.5- fold to 37.5-fold. Notably, there is an 81-fold increase in the promoter activity if we compare the relative activity of the variant (-1925/+52) to that of (-1442/+52). When we keep deleting the -97/+52 sequence, the activity decreases stepwise to 37.5-fold (-77/+52), 8.54-fold (- 37/+52), and 0.06-fold (-17/+52).

**Figure 1.**
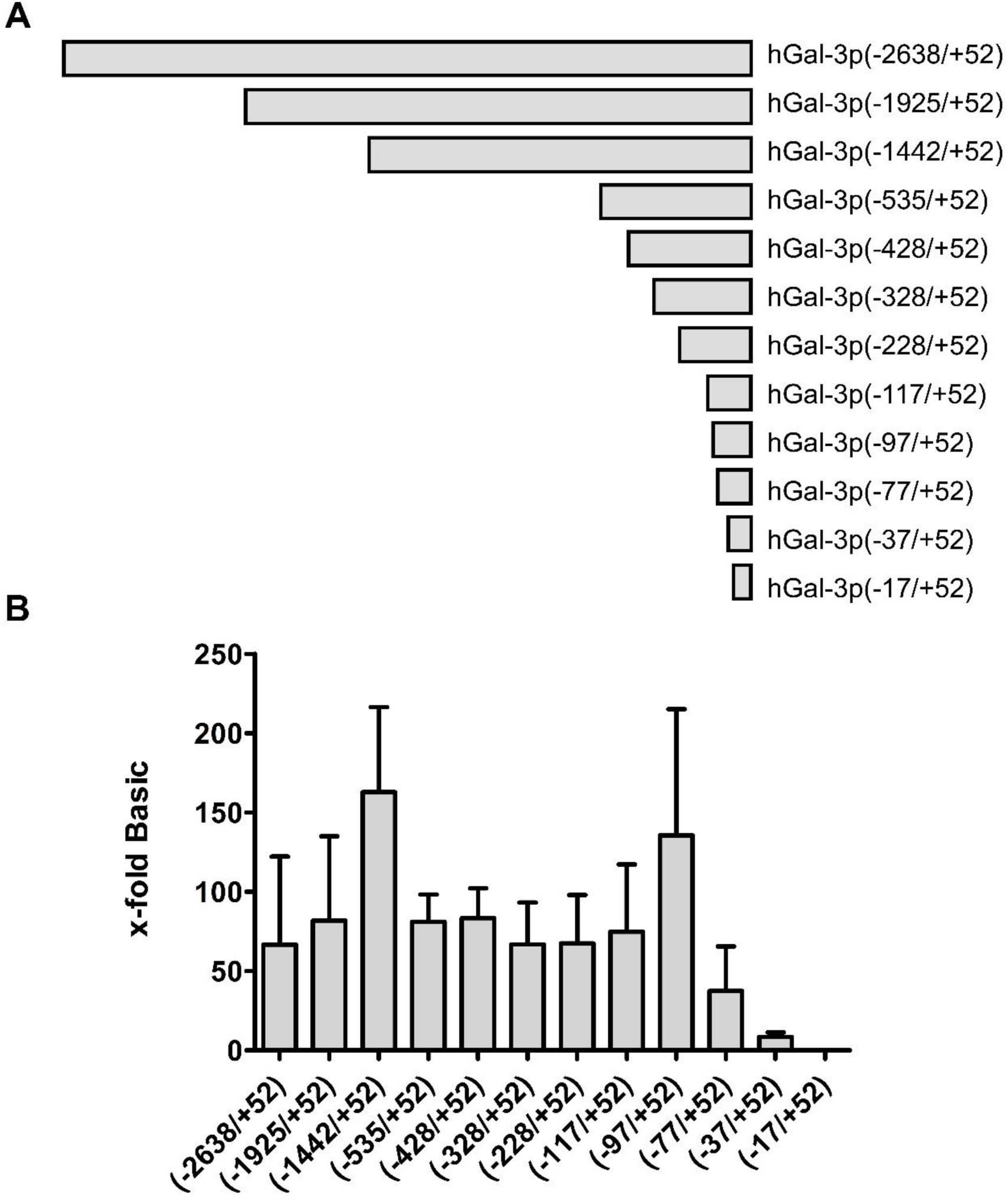
(A) The LGALS3 full-length promoter (-2638/+52) and the deletions (-1925/+52), (-1442/+52), (-535/+52), (-428/+52), (-328/+52), (-228/+52), (-117/+52), (-97/+52), (-77/+52), (-37/+52), (-17/+52) are shown in a diagram drawn to scale. Positions are numbered according to the distance to the transcription start site (TSS), where negative means upstream and positive means downstream of the TSS. (B) Relative activity of LGALS3 promoter sequence deletions in SW1353 cells. The activity of these deletions is shown as an x-fold increase relative to cells transfected with the promoter-less vector pGL4.20 (Basic). The graph shows the results’ mean ± SD of at least four biological replicates, each including three technical replicates, with error bars representing the standard deviation. *p < 0.05.

To investigate whether these observed activity levels correspond to a conserved sequence across species, we aligned the region 2690bp upstream of the *LGALS3* TSS of selected species with the human *LGALS3* promoter sequence (-2638/+52). The alignment revealed conserved motifs within this upstream region, mainly in primates (positions -600 to 0 relative to human TSS) and rodents (positions -2200 to -1200 relative to human TSS), suggesting that this promoter is of evolutionary significance (Supplementary Figures 1 and 2). However, we could not identify conserved regions among other species, e.g., chicken, by using a minimum conservation threshold of 50%.

### 3.2. Effects of SOX proteins on the activity of the *LGALS3* promoter

We first performed a qPCR analysis to characterize the cell line SW1353 regarding the expression of *SOX* mRNA. As mentioned, SOX proteins are transcription factors that are dysregulated within OA and could be involved in the regulation of galectins. The mRNA levels of the chosen SOX genes (*SOX2, SOX4, SOX5, SOX6,* and *SOX9*) relative to that of GAPDH ranged from 10^-3^ to 10^-4^ (Supplementary Figure 3). *SOX2* and *SOX9* were the genes with the lowest relative expression level in the SW1353 cells, yet the Kruskal-Wallis test did not find the difference within the group of SOX genes to be significant.

Consecutively, by co-transfecting SW1353 cells with 50 ng of a reporter construct containing the full-length *LGALS3* promoter, pGL4-hGal3p(-2638/+52), and 30 ng of plasmids encoding different SOX genes (pcDNA3.1(+)-SOX2, pcDNA3.1(+)-SOX4, pcDNA3.1(+)-SOX5, pcDNA3.1(+)-SOX6, pcDNA3.1(+)-SOX9), we investigated quantitatively their effect on the promoter activity. The vector pcDNA3.1(+), without any SOX encoding sequence, was used as a control. The relative luciferase activities of the full-length promoter in the presence of the SOX proteins were calculated as a percentage referred to the 100% value of the control (Figure 2). To confirm that transfections with SOX-encoding plasmids were efficient and the SOX- encoding plasmids caused changes in promoter activity, an additional 24-well plate was processed in the same way and used for qPCR analysis of the SOX genes’ mRNA expression (data not shown). The *SOX2*-containing vector caused repression of *LGALS3* promoter activity by 80.1 %, and the *SOX9*-containing vector by 48.3 %. In contrast, the *SOX4*-, *SOX5*-, and *SOX6*-containing vectors increased promoter activity by 23.9 %, 36.5 %, and 22.5 %, respectively. The only statistically significant difference found was between *SOX2* and *SOX5* (p < 0.05).

**Figure 2.**
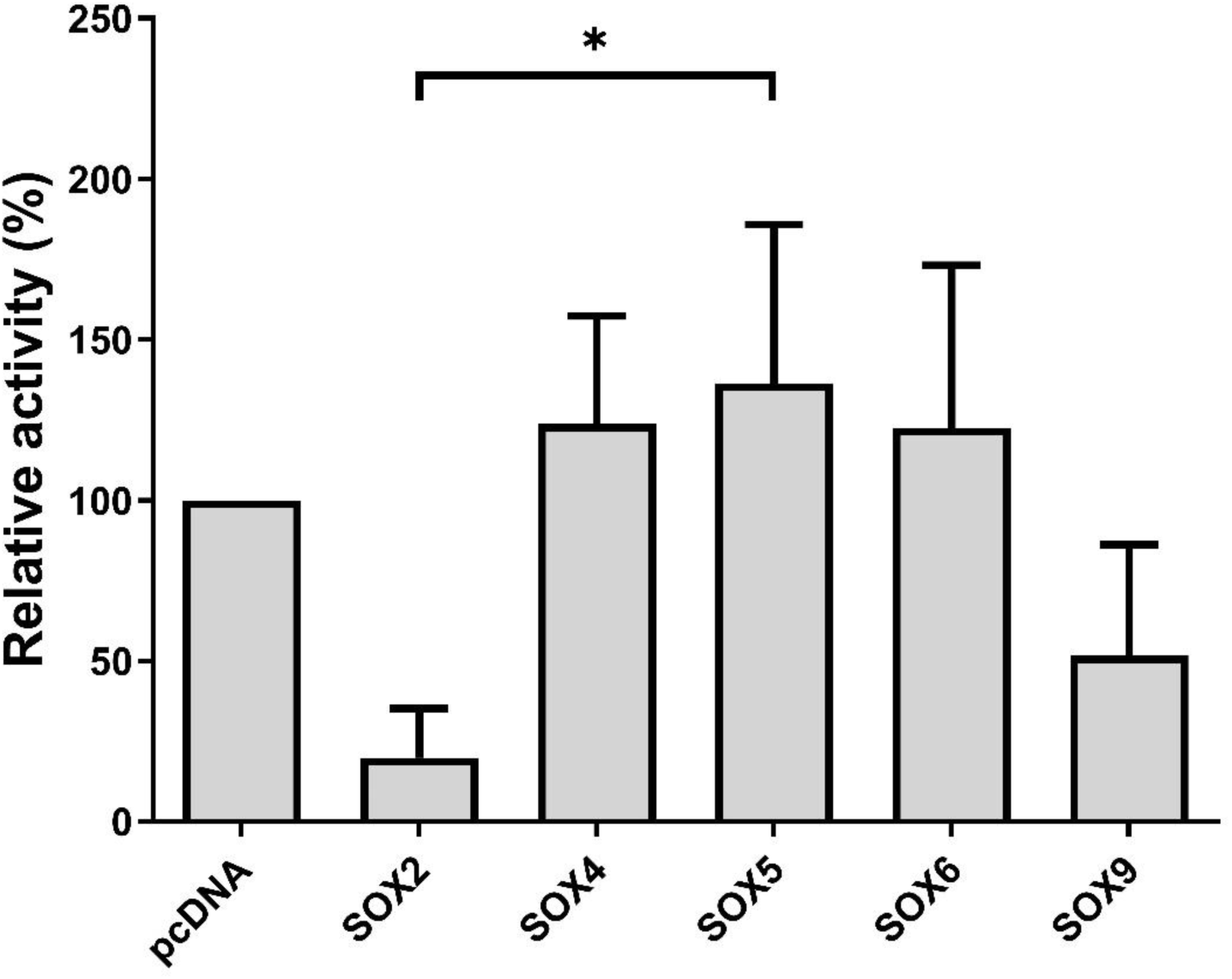
Relative activity of the LGALS3 full-length promoter (-2638/+52) in SW1353 cells when co- transfected with 30 ng of different SOX-containing plasmids (pcDNA3.1(+)-SOX2, -SOX4, -SOX5, - SOX6 and -SOX9). The relative activity of the promoter in the different conditions is calculated as a ratio to the promoter activity of the cells transfected with the vector pcDNA3.1(+) lacking the respective SOX. The graph shows results’ mean relative activity ± SD from four technical replicates, each including three biological replicates, with error bars representing the standard deviation. *p < 0.05.

### 3.3. *SOX2* and *SOX9* regulate the *LGALS3* promoter activity in a dose-dependent manner

To determine dose dependency of *LGALS3* promoter activation by SOX proteins, we repeated the co-transfection experiment mentioned in the previous section with increasing amounts of pcDNA3.1(+)-SOX2, pcDNA3.1(+)-SOX4, pcDNA3.1(+)-SOX5, pcDNA3.1(+)-SOX6 or pcDNA3.1(+)-SOX9 plasmid DNA (0.1, 1, 10, 100 ng). With increasing amounts of *SOX2-* and *SOX9*-encoding plasmids, we observed a stepwise decrease of *LGALS3* promoter activity. A statistically significant difference in relative activity was detected between samples with the lowest (0.1 ng) and highest (100 ng) plasmid amounts for both *SOX2*- and *SOX9*-encoding constructs (Figure 3). By transfecting cells with 100 ng of the respective plasmids, we achieved a repression of the *LGALS3* promoter of 97.2 % with *SOX2* and 92.6 % for *SOX9*. These findings demonstrate that overexpression of either *SOX2* or *SOX9* alone can suppress *LGALS3* promoter activity in SW1353 cells.

**Figure 3.**
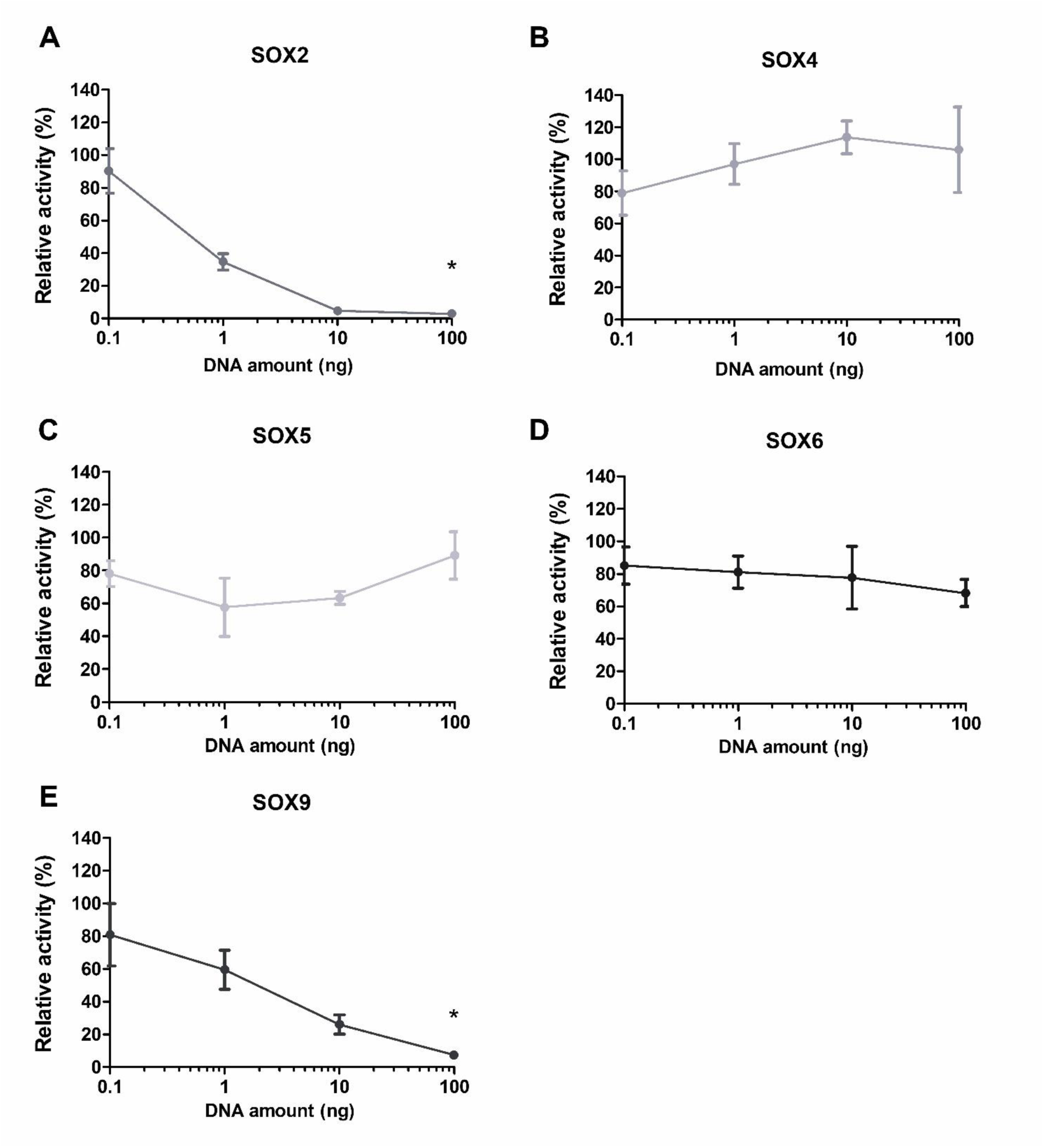
Relative activity of the LGALS3 full-length promoter (-2638/+52) in SW1353 cells when titrated with different amounts of SOX-containing plasmids. (A) pcDNA3.1(+)-SOX2, (B) pcDNA3.1(+)- SOX4, (C) pcDNA3.1(+)-SOX5, (D) pcDNA3.1(+)-SOX6, (E) pcDNA3.1(+)-SOX9. The amounts of DNA used were: 0.1 ng, 1 ng, 10 ng, 100 ng. For each condition, a negative control without the SOX-encoding plasmid was included. The x-axis is plotted on a logarithmic scale. The relative activity of the promoter in the different conditions is calculated as a ratio to the promoter activity of the cells transfected with the vector pcDNA3.1(+) lacking the respective SOX. The graph shows results’ mean relative activity ± SD from three biological replicates, including three technical replicates, with error bars representing the standard deviation. *p < 0.05.

Transfection with varying amounts of pcDNA3.1(+)-SOX4, pcDNA3.1(+)-SOX5, and pcDNA3.1(+)-SOX6 did not significantly change *LGALS3* promoter activity, e.g., in the case of pcDNA3.1(+)-SOX6 transfection, the promoter activity level remained stable between 68.1 % and 85.1 %. Notably, the promoter activity increased when transfected with 0.1 ng to 10 ng of pcDNA3.1(+)-SOX4, but not at 100 ng.

### 3.4. *In silic*o analysis of the *LGALS3* promoter for transcription factor binding sites (TFBS) revealed multiple potential SOX binding sites

An *in silico* analysis using the JASPAR database was performed to find potential SOX-binding sites within the human *LGALS3* promoter region. This analysis aimed to determine whether the observed SOX-mediated effects on *LGALS3* expression could be due to a potential binding within the *LGALS3* promoter sequence. SRY matrices were also included, as SRY and other SOX proteins bind to DNA through their HMG-box domains and are part of the same family: the SOX (SRY-related HMG-box) family. JASPAR predicted 655 SOX-binding sites within the full-length (-2638/+52) *LGALS3* promoter at a threshold of 80%. The 20 results of SOX TBFS with the highest relative score and their relative positions within the promoter sequence are presented in Supplementary Table S5. A region spanning from -505 to +52 relative to the TSS is shown with SOX binding sites at a 70% threshold in Figure 4. We could observe a higher density of SOX TFBS from the -505 to -325 region than in the -325 to +52 region.

**Figure 4.**
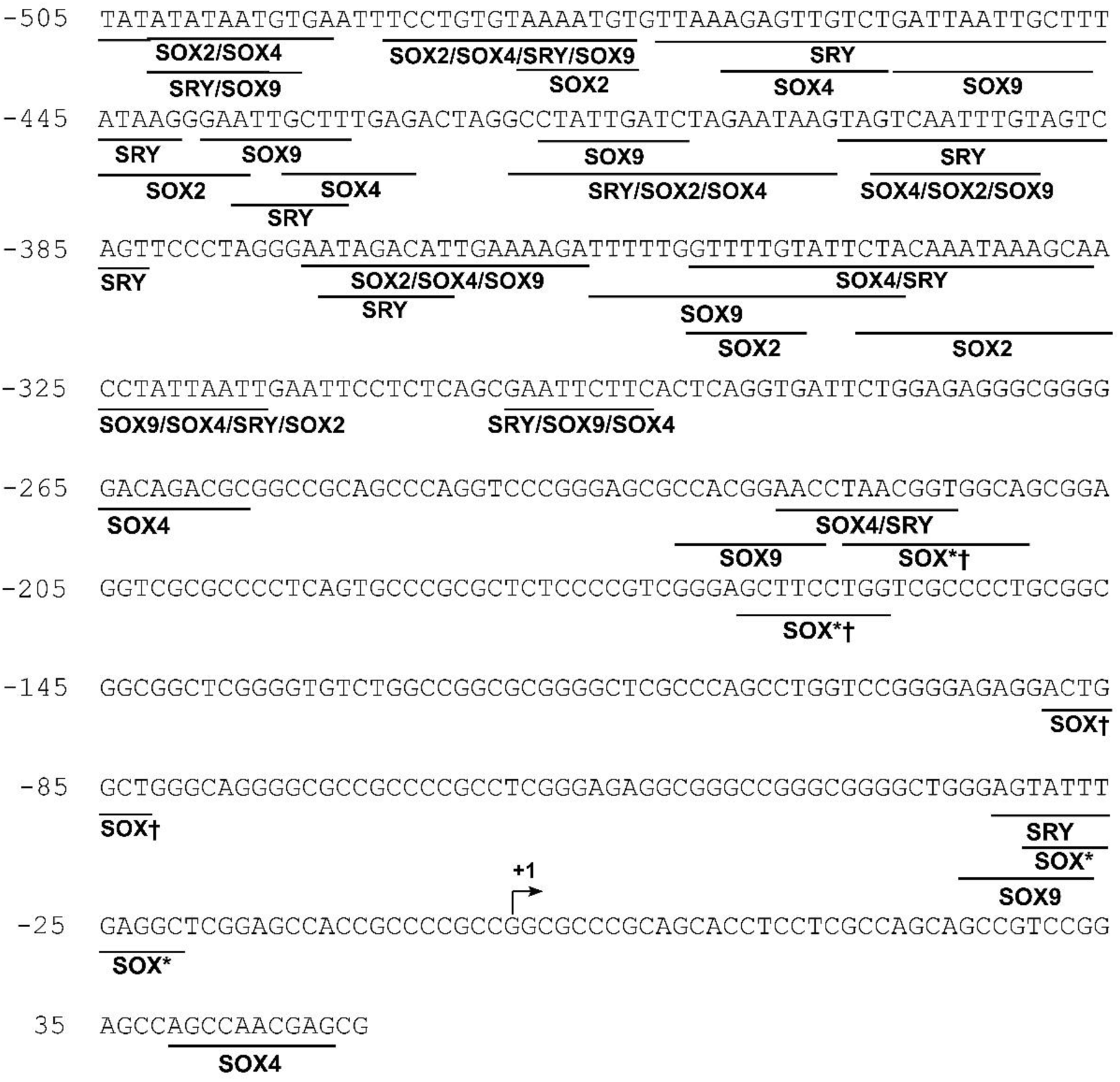
Human LGALS3 promoter (-505/+52) with potential SOX binding sites. Information on the binding sites was obtained from JASPAR, setting the threshold at 70%. TSS is indicated with an arrow.

SOX9 (Matrix ID MA0077.1.SOX9) could potentially bind to the *LGALS3* promoter on multiple sites, for instance, on the sense strand (+) at positions -418 to -410 relative to the TSS, with a high relative score (0.92, Figure 4). For SOX2, multiple putative binding sites were also found (Matrix ID MA0143.5.SOX2) predicted with high relative scores of up to 0.88 (result not shown) to bind to the *LGALS3* promoter on the sense strand (+) at positions -1462 to -1456 relative to the TSS, and positions -1133 to -1127 relative to the TSS on the antisense strand (-). SOX4 (Matrix IDs MA0867.3 and MA0867.2.SOX4) binding sites had high relative scores (up to 0.95) at positions -1833 to -1926 relative to the TSS (Supplementary Table S5).

The analysis also found some SOX proteins that were not the object of this study. However, it is worth mentioning that a binding motif for SOX10 (Matrix IDs MA0442.3 and MA0442.1) was found at position -1698 to -1693 relative to the TSS, consisting of the complementary sequences 3’-ACAAAG-5’ and 5’-CTTTGT-3’ (Supplementary Table S5). Additionally, we found multiple potential SOX15 (Matrix ID MA1152.2) binding sites (relative scores ranging from 0.93 to 0.98) at various positions, including -1833 to -1827 and -2243 to -2237 relative to the TSS (Supplementary Table S5). These potential binding sites suggest that SOX proteins could bind to the *LGALS3* promoter, possibly impacting its regulation. In addition to TFBS for SOX proteins, we found several motifs for other TFs (e.g., KLFs), presented in Supplementary Table S6.

### 3.5. Effect of *SOX2* and *SOX9* on promoter deletion constructs

To explore which sequence stretches are responsible for the SOX2- and SOX9-mediated effects, we examined the impact of pcDNA3.1(+)-SOX2 and pcDNA3.1(+)-SOX9 on *LGALS3* promoter deletion constructs (Figure 5). Luciferase assays showed that *SOX2* exerted similar repression levels on the activity of each deletion variant, with a residual promoter activity between 6.3 ± 1.9 % and 9.3 ± 2.8 % (Figure 5A). For *SOX9* (Figure 5B), the level of repression was slightly different between the analysed constructs, but still, a relative activity of 25.5 ± 4.0 % of the proximal hGal-3p(-97/+52) promoter region can be observed. Promoter activity was highest for the hGal-3p(-1442/+52) construct (41.3 ± 8.6 %) and lowest for the full-length *LGALS3* promoter (-2638/+52) (18.4 ± 6.2 %) (Figure 5B). When comparing the full-length promoter (-2638/+52) with the following deletion construct (-1442/+52) in Figure 5B, activity increased, but diminished when measuring shorter promoter constructs. This analysis allowed us to narrow down the specific sequence region responsible for the SOX-mediated repression to the -97/+52 region.

**Figure 5.**
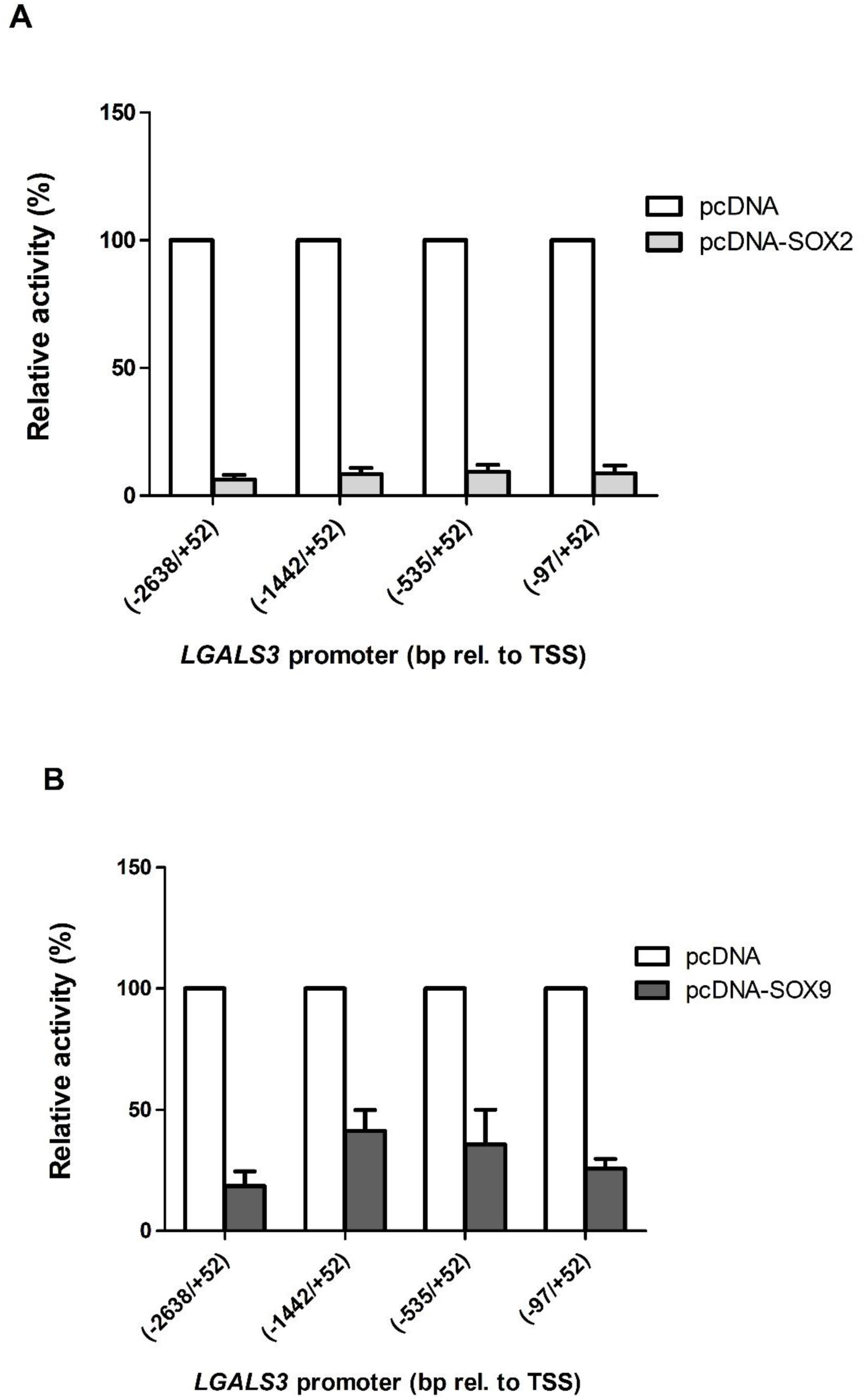
Relative activity of LGALS3 promoter deletion variants in SW1353 cells when co-transfected for 24 h with SOX-protein-containing plasmids: (A) SOX2 or (B) SOX9. The graph shows results’ mean ± SD from at least three biological replicates, including three technical replicates, with error bars representing the standard deviation. The white bars represent the negative control (empty pcDNA3.1(+)). (A) The light grey bars represent SOX2. (B) The dark grey bars represent SOX9.

### 3.6. SOX9 can directly bind to the *LGALS3* promoter

To assess whether the observed repressive effect of SOX9 on the *LGALS3* promoter activity was due to direct binding of the promoter, we performed a chromatin immunoprecipitation using HaloTag plasmid pHTN-SOX9. We focused on SOX9 because it is a key regulator in cartilage and chondrogenesis, and its transcription activates genes for many cartilage-specific components and regulatory factors. As a positive control, we used the *ADAMTS*-4 (disintegrin and metalloprotease with thrombospondin-1-like domains-4) promoter, which is known to interact with the SOX9 protein [30]. In our analysis, SOX9 showed an efficient binding to the *ADAMTS-4* promoter area, leading to a 25-fold enrichment in our CHIP-qPCR compared to the untransfected cells (control, Figure 6A). For *LGALS3,* we obtained a 17-fold increase for the sequence between -93 and +49 relative to the TSS with cells transfected, proving the binding of SOX9 within this sequence stretch. Figure 6B shows the studied sequence stretch, with a potential SOX9 binding site highlighted in a box, and in Figure 6C, a canonical SOX9 binding motif is shown.

**Figure 6.**
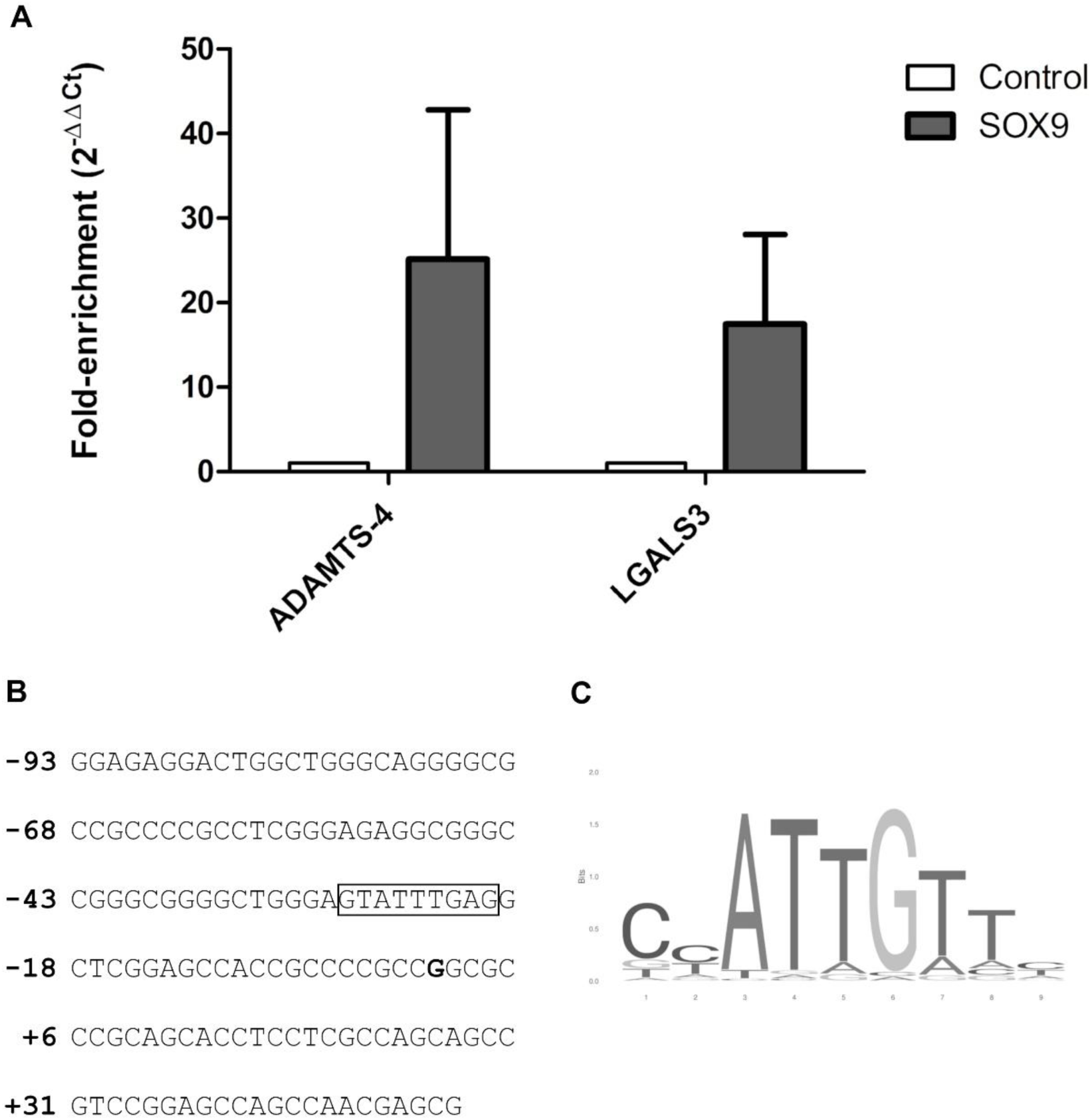
Fold enrichment of specific promoter binding of HaloTag-SOX9 and hGal3p(-93/+49) in vitro. (A) HaloChIP experiments were performed using an in vitro system with recombinant HaloTag-SOX9 or a negative control (-). DNA bound to SOX9 was isolated using the HaloCHIPTM system (Promega), and DNA was amplified via quantitative PCR. ADAMTS-4 is a known target of SOX9, which was used as a positive control. The graph shows results’ mean ± SD from two biological replicates, with error bars representing the standard deviation. The white bars represent the negative control (HaloTag without SOX9 attached). The dark grey bars represent HaloTag-SOX9. (B) The sequence stretch studied is shown, with a potential SOX9 binding site shown in a box. (C) A canonical SOX9 binding motif is shown.

## 4. Discussion

Galectin-3 is a multifunctional β-galactoside-binding lectin implicated in various cellular processes [4,10,31–46]. Elevated galectin-3 levels occur in several pathological conditions, including OA [10]. We previously reported that the upregulation of galectin-3 in OA cartilage is associated with increased tissue degeneration [10,47]. However, the molecular mechanisms underlying this upregulation remain poorly understood. In particular, the transcriptional regulation of the *LGALS3* gene expression and promoter activity has not been investigated in the chondrocyte context.

The first aim of this study was to pinpoint the essential sequence regions that regulate the *LGALS3* promoter in the SW1353 chondrocyte model. A series of deletion constructs, from 17 bp to 2638 bp upstream of the TSS, enabled detailed promoter sequence analysis. Notably, the level of promoter activity remained unchanged despite the removal of up to 2541 bp, indicating that the minimal promoter region from -97 to +52 relative to the TSS is sufficient to drive transcription. This finding means that a fragment of 149 bp constitutes an active promoter containing the key regulatory elements, activating transcription of the *LGALS3* gene in this cellular context.

We further showed an increase in promoter activity of 81.1 % if we compare the relative activity of construct hGal3p(-1925/+52) to that of hGal3p(-1442/+52). This 500 bp region will probably contain repressive regulatory elements, whose removal leads to enhanced transcriptional activity. In contrast, comparing the promoter activity of the hGal3p(-97/+52) construct to that of the hGal3p(-77/+52), we observed a marked decrease in activity, indicating that the 20 bp segment between -97 and -77 contains sequences essential in activating gene expression. Deletion of this region likely impaired the binding of transcriptional activators or disrupted other regulatory features necessary for full promoter function.

Using JASPAR *in silico* analysis, we identified transcription factor binding motifs (sites) for SP2, PATZ1, KLF15 (Supplementary Table S6), and others within the -97/+52 minimal *LGALS3* promoter region, which may contribute to *LGALS3* transcriptional activation. *In silico* analysis of the full-length promoter identified 655 potential SOX-binding sites at a threshold of 80%, suggesting a possible regulatory involvement of SOX proteins in *LGALS3* gene expression.

SOX proteins are essential regulators throughout chondrogenesis [16,48–51], whereas in OA, several SOX proteins are dysregulated and contribute to disease progression [14,17,18,30]. The SOX trio (SOX5, SOX6, and SOX9), which is essential for cartilage development and maintenance [16,48], shows reduced expression in osteoarthritic cartilage, suggesting a loss of chondrocyte identity and function [17], while SOX4 directly induces expression of catabolic enzymes such as ADAMTS-4 and ADAMTS-5 [18]. Likewise, SOX9 has been implicated in regulating cartilage degeneration associated with ADAMTS activity in early OA [30].

Therefore, we hypothesised that SOX proteins (SOX2, SOX4, SOX5, SOX6, and SOX9) are involved in regulating the *LGALS3* promoter in OA. This hypothesis is primarily supported by our results, which originated from a series of transfections using SOX-expressing plasmids. We provided evidence for a regulatory link between SOX factors and *LGALS3*. Specifically, we showed that *SOX2* and *SOX9* are repressors of the *LGALS3* promoter (Figures 2 and 3). Titration experiments with SOX-expressing plasmids ranging from 0.1 to 100 ng revealed a dose-dependent decrease in *LGALS3* promoter activity.

While SOX2 and SOX9 significantly reduced *LGALS3* promoter activity in luciferase reporter assays, SOX5 and SOX6 did not show any notable effect. In contrast, SOX4 showed a dose-dependent increase of the promoter activity at lower concentrations (0.1–10 ng), which was not maintained at higher levels (100 ng). Since *SOX2* and *SOX9* showed the most pronounced effects, we tested the pcDNA3.1(+)-SOX2 and SOX9 constructs with different *LGALS3* promoter deletion variants to narrow the specific regions crucial for repression. Our results indicate that *SOX2*-mediated repression occurs irrespective of the promoter region deleted, with a range of residual promoter activity of 6.3 % to 9.3 % (Figure 5A). In contrast, SOX9*-* induced repression of the promoter activity was weaker for the deletion construct (-1442/+52) than for the full-length promoter, which could indicate the presence of the repressor SOX9 TFBS within the -2638bp to -1442bp area, whose deletion led to an increase in the activity. From the (-1442/+52) to the (-97/+52) construct, the activity decreased steadily but not significantly, meaning that activating elements could be removed. Whether these elements include SOX9 TFBS remains unknown. Our findings revealed that SOX2 and SOX9 are transcriptional repressors of *LGALS3* expression. Remarkably, SOX9 - a key regulator of chondrogenesis - is downregulated in OA cartilage [52], while we showed that galectin-3 expression in OA cartilage is increased [10]. The reduced SOX9 expression in OA chondrocytes may relieve repression of the *LGALS3* promoter, potentially contributing to the observed upregulation of galectin-3 in OA [10,47].

Interestingly, galectins have also been shown to regulate SOX transcription factors. Galectin- 1 has been reported to regulate the expression of SOX9 in colorectal cancer models [19]. Furthermore, galectin-3 has been implicated in the modulation of SOX2 expression during osteogenic differentiation of human periodontal ligament stem cells [53]. These findings suggest an engagement of galectins and SOX proteins in a greater regulatory feedback loop. Song et al. (2005) [43] showed that galectin-3 could form a complex with AP-1 and DNA, as demonstrated by chromatin immunoprecipitation and pull-down. Since AP-1 is also known to interact with SOX9 [51], this could further indicate how different proteins interact in this context, highlighting the critical role of SOX proteins in galectin expression.

Zhang et al. (2015) [30] reported SOX9 as an activator and repressor, describing that “SOX9 repressed ADAMTS’ expression and promoted COL2A1, ACAN, and COMP expression in human chondrocytes”. Our findings support that SOX9 represses *LGALS3* promoter activity. To substantiate this claim, we provide evidence that SOX9 can bind directly to a respective transcription factor site within the -93/+49 region of the *LGALS3* promoter, with a 17-fold enrichment observed in our HALO-ChIP assay compared to untransfected cells (Figure 6A). This result is the first proof of direct SOX9 binding to the *LGALS3* promoter, identifying a potentially critical regulatory mechanism for galectin-3 dysregulation in OA.

However, despite direct binding to the promoter, we cannot exclude that SOX9 may also repress the *LGALS3* promoter through other mechanisms, such as inhibition of the Wnt/β- catenin signalling pathway. TFBS analysis using JASPAR identified several high-scoring TCF binding sites in the *LGALS3* promoter (with a 70% threshold), indicating a potential role for regulation through the Wnt/β-catenin signalling. As shown in Supplementary Figure 5, we observed TCF3, 4, and 12 binding sites scattered across the (-505/+52) region, and TCFL5 (at positions (-205/-195)) and (-191/-181)), TCFL7 (at position (-27/-20)), and TCFL21 (at position (+26/+34)). SOX9 possesses an inhibitory activity through the Wnt/β-catenin pathway [49,54] by direct competition with TCF/Lef for β-catenin binding, leading to its degradation [54,55]. We could hypothesise that this competitive binding is responsible for SOX9-mediated repression of *LGALS3* expression. Thus, the inhibitory effect of SOX9 on *LGALS3* transcription could result from direct promoter binding or indirect regulation via β-catenin degradation, potentially involving Wnt signalling.

Since our overall aim was to investigate the regulation of *LGALS3* in OA, the question arose as to what extent the SW1353 chondrosarcoma cells can be a model for OA chondrocytes. Therefore, we analysed the mRNA expression levels of *LGALS3* and the SOX transcription factors of interest in OA primary chondrocytes (Supplementary Figure 6A) and compared them with those in the SW1353 cells (Supplementary Figure 6B). We observed that *LGALS3, SOX4, SOX5* and *SOX9* were expressed at higher levels than in SW1353 cells (Supplementary Figure 6B). Although, as demonstrated in this study, we can establish basic principles of *LGALS3* regulation by SOX proteins in SW1353 cells. Future studies should investigate these mechanisms in OA primary chondrocytes to validate the SOX proteins controlling the *LGALS3* promoter in the OA setting.

In conclusion, we have identified the minimal active promoter sequence of human *LGALS3* and quantified the effect of *SOX2*, *SOX4*, *SOX5*, *SOX6*, and *SOX9* on its activity. We observed stable promoter activity until region -2638/-97 was deleted, highlighting the importance of the -97/+52 region. We demonstrated a repressive effect of *SOX2* and *SOX9* on the human *LGALS3* promoter activity and revealed SOX9 binding to the -97/+52 region. Furthermore, we propose a feedback regulatory loop between *SOX2* and *SOX9*. Our study opens up new possibilities for advancing our understanding of galectin regulation in the context of OA and other inflammation-associated diseases.

## Ethical approval

The use of clinical specimens for this study was approved by the ethics committees of the Medical University of Vienna (EK-No.: 1922/2017 and 1555/2019).

## Declaration of Generative AI and AI-assisted technologies in the writing process

During the preparation of this manuscript, the authors did not use any AI.

## Data availability

Data will be made available on request.

## CRediT authorship contribution statement

Blanca Alba: Investigation, Visualization, Formal analysis, Validation, Writing – Original Draft, Writing – Review & Editing.

Herbert Kaltner: Supervision, Writing – Review & Editing. Stefan Toegel: Resources, Writing – Review & Editing.

Sebastian Schmidt: Conceptualization, Investigation, Visualization, Project Administration, Data Curation, Validation, Writing – Original Draft, Writing – Review & Editing.

## Acknowledgements

We gratefully acknowledge the technical assistance by Christian Overdiek, Martin Vogel, and Melanie Cezanne.

