## Supplementary Material for "SOX2 and SOX9 as Transcriptional Regulators of human Galectin-3 in SW1353 Cells: Potential Implications for Osteoarthritis"

Blanca Alba, Herbert Kaltner^,^ Stefan Toegel & Sebastian Schmidt*

**SUPPLEMENTARY MATERIAL**

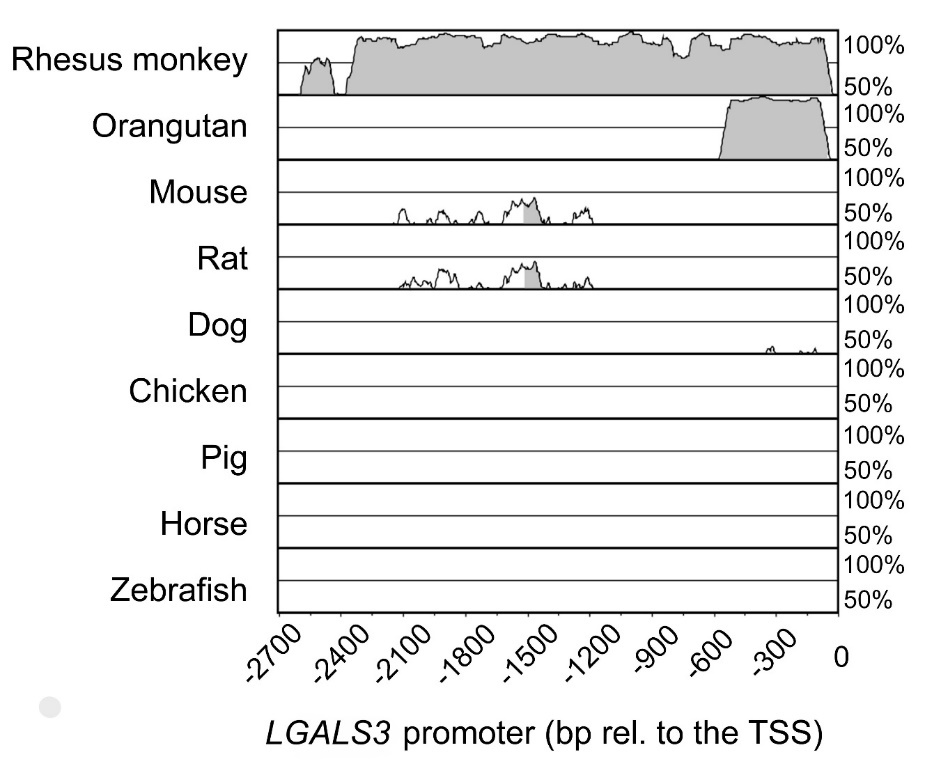

Supplementary Figure 1. Alignment of the *LGALS3* promoter of 9 species to the human sequence (Homo sapiens, Gene ID: 3958) using the mVISTA software. The species used were: Rhesus monkey (Macaca mulatta, Gene ID: 715848), orang-utan (Pongo abelii, Gene ID: 100453870), mouse (Mus musculus, Gene ID: 16854), rat (Rattus norvegicus, Gene ID: 25664), dog (Canis lupus familiaris, Gene ID: 403832), chicken (Gallus gallus, Gene ID: 373917), pig (Sus scrofa, Gene ID: 100038033), horse (Equus ferus caballus, Gene ID: 100063828) and zebrafish (Danio rerio, Gene ID: 100006084). Grade of conservation in percent is given on the right axis. The relative position relative to the TSS is given below.

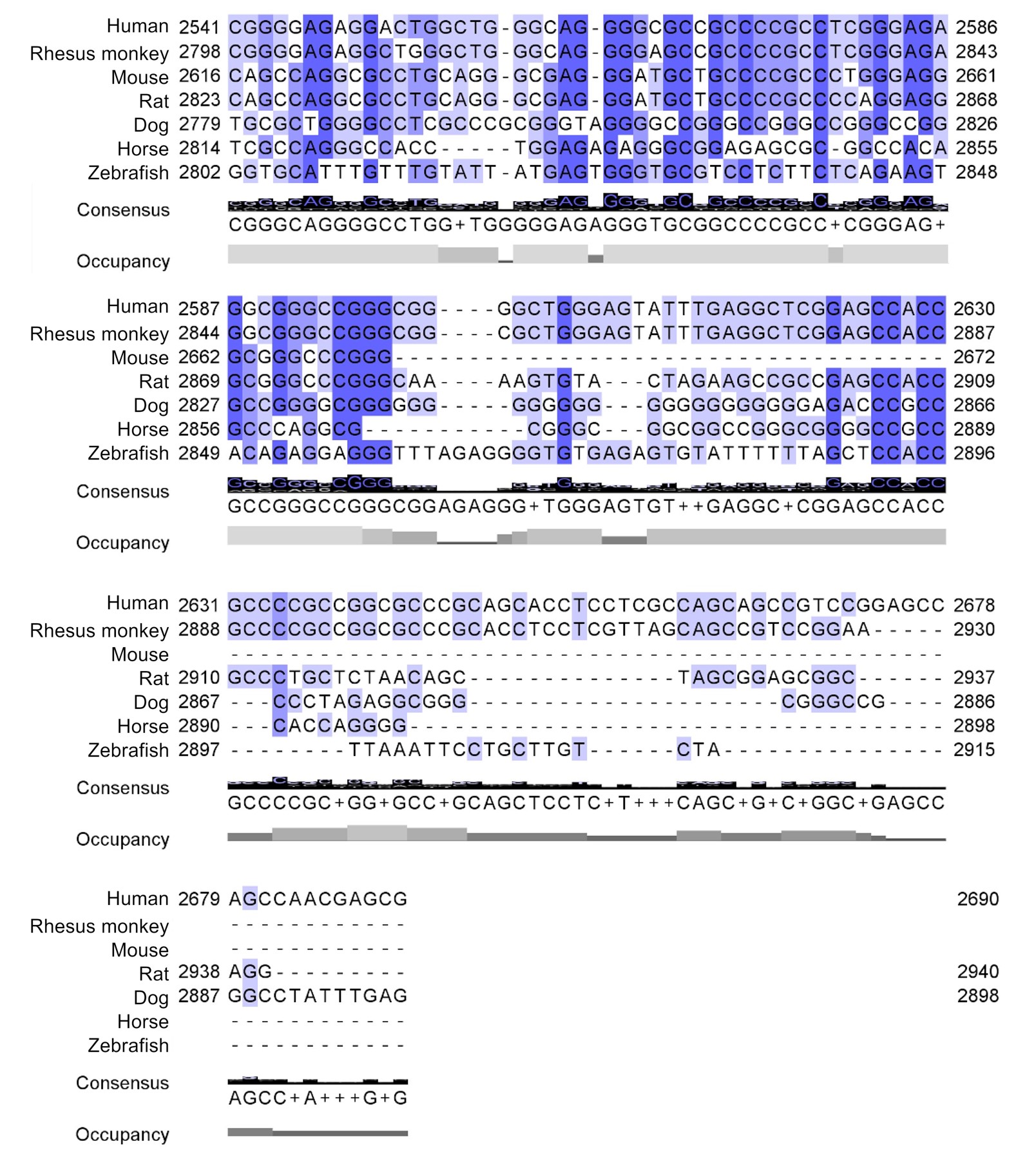

Supplementary Figure 2. Alignment of the -97/+52 region of *LGALS3* promoter of 9 species to the human sequence (Homo sapiens, Gene ID: 3958) using the Jalview software. The species used were: Rhesus monkey (Macaca mulatta, Gene ID: 715848), orang-utan (Pongo abelii, Gene ID: 100453870), mouse (Mus musculus, Gene ID: 16854), rat (Rattus norvegicus, Gene ID: 25664), dog (Canis lupus familiaris, Gene ID: 403832), chicken (Gallus gallus, Gene ID: 373917), pig (Sus scrofa, Gene ID: 100038033), horse (Equus ferus caballus, Gene ID: 100063828) and zebrafish (Danio rerio, Gene ID: 100006084). The consensus sequence for all species is given below.

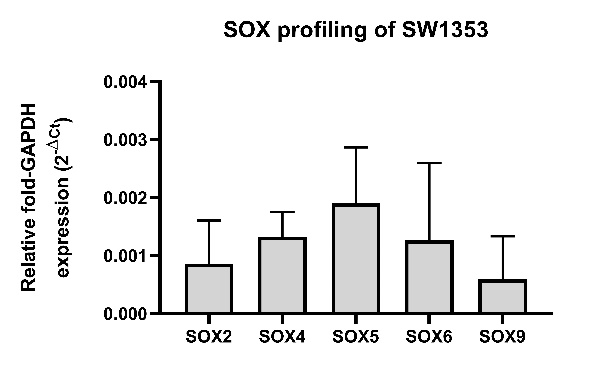

Supplementary Figure 3. Quantitative PCR analysis of SOX genes (*SOX2*, *SOX4*, *SOX5*, *SOX6* and *SOX9*) mRNA expression in SW1353 cells. The graph shows results’ mean ± SD from three biological replicates, each including three technical replicates, with error bars representing the standard deviation. RNA was extracted from a SW1353 cell pellet, reverse transcribed into cDNA and analysed by qPCR in order to measure the relative gene expression of the SOX genes of choice: *SOX2*, *SOX4*, *SOX5*, *SOX6* and *SOX9*.

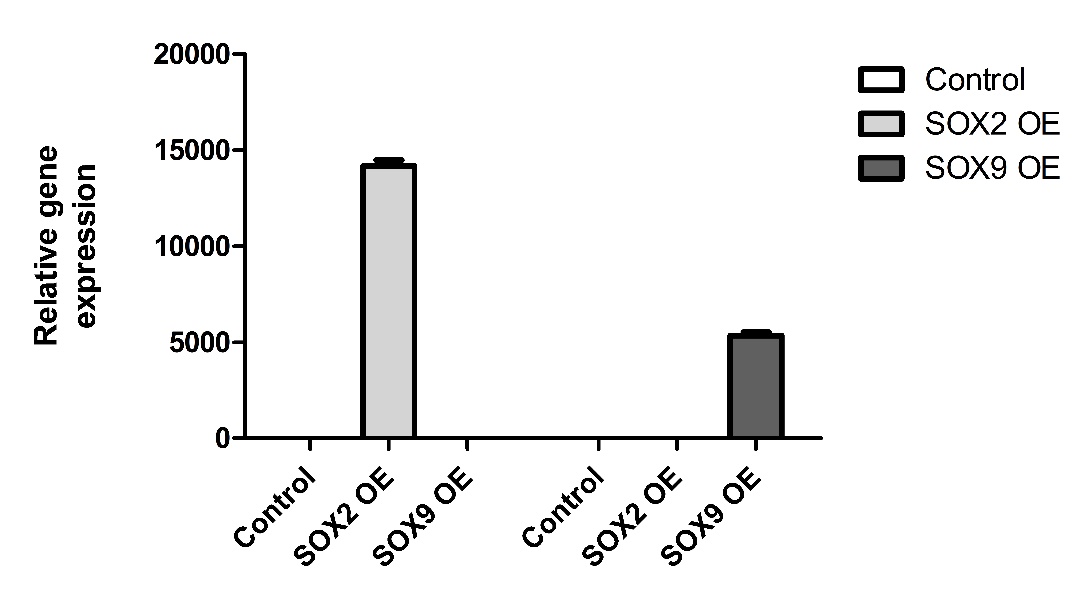

Supplementary Figure 4. Quantitative PCR analysis of SOX genes (*SOX2* and *SOX9*) mRNA expression in SW1353 cells using the Pfaffl method, 48h after transfecting with pcDNA-SOX2 and pcDNA-SOX9. The graph shows results’ mean from three technical replicates, with error bars representing the standard deviation. RNA was extracted from a SW1353 cell pellet, reverse transcribed into cDNA and analysed by qPCR in order to measure the relative gene expression of the *SOX2* and *SOX9*.

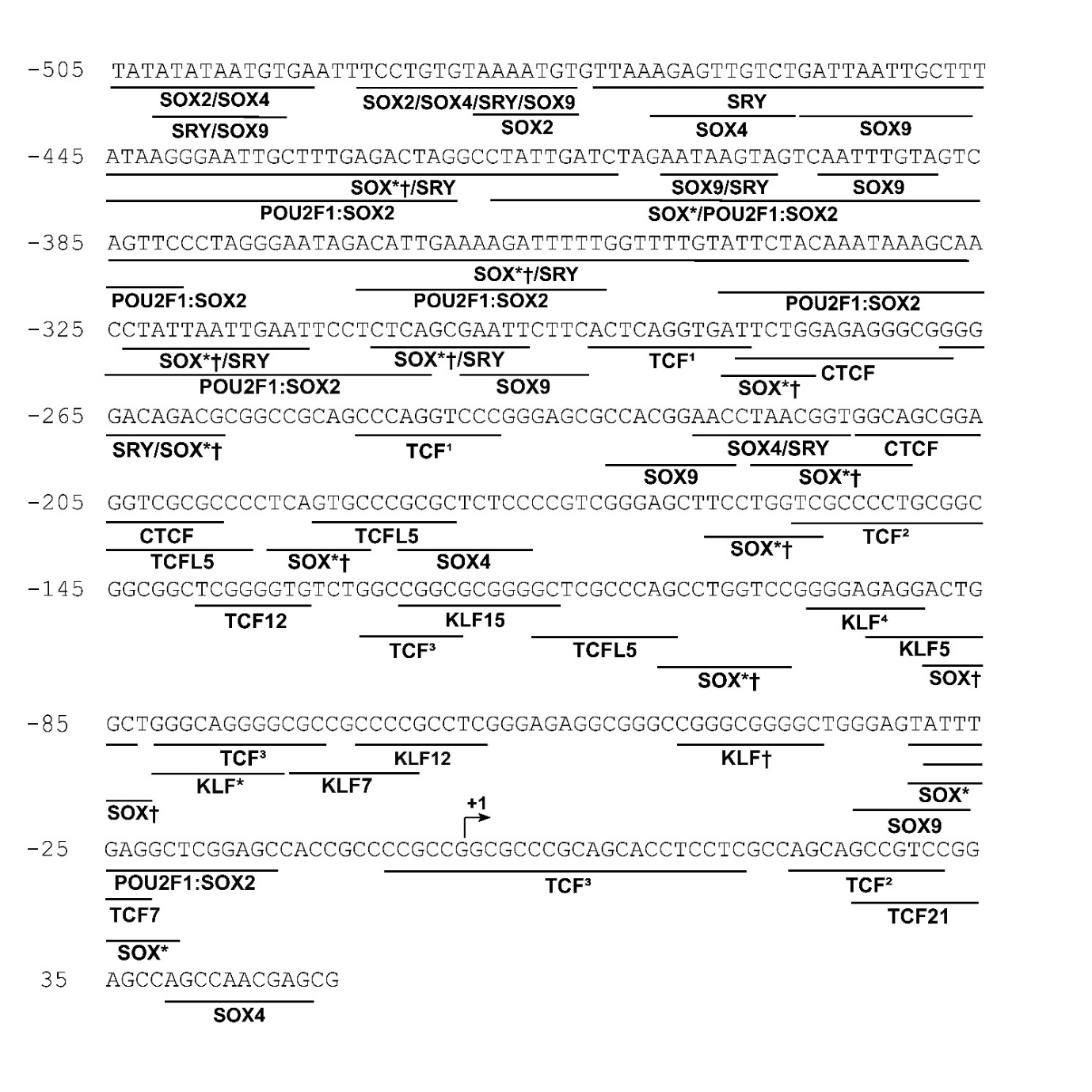
Supplementary Figure 5. Human *LGALS3* promoter (-505/+52) with SOX, TCF and KLF binding sites. Information of the binding sites was obtained from JASPAR, setting the threshold at 70%. Legend is as follows: TCF1 for TCF3 and 4. TCF2 for TCF3 and 12. KLF1 for KLF1, 4, and 7. KLF† for KLF10, 12, and 14. KLF* for KLF1, 2, 4, 7 and 15. SOX* for SOX2, 4 and SRY. SOX† for SOX9 and SRY. SOX†* for SOX2, 4 and 9. The TSS is indicated by an arrow.

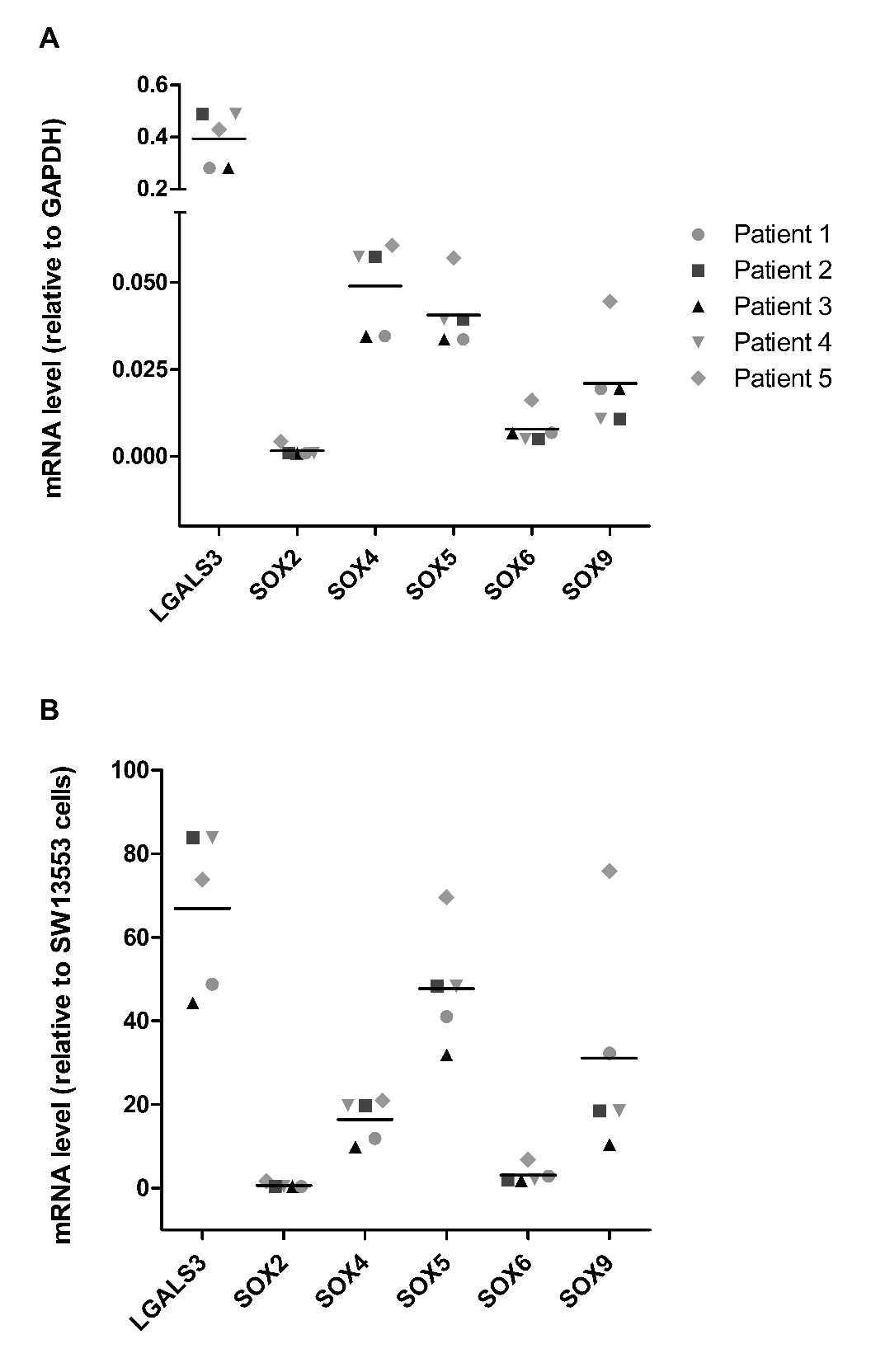

Supplementary Figure 6. Quantitative PCR analysis of *LGALS3* and *SOX* mRNA expression levels from patient material and SW1353 cells. (A) mRNA level (relative to GAPDH), with the expression level of GAPDH set to 1, serving as the reference for comparison. (B) mRNA level (relative to SW13553 cells), with the expression level in SW1353 set to 1, serving as the reference for comparison. The results shown are from two technical replicates performed with five biological replicates.

Supplementary Table S1. Vectors.^[1]^

| Vector name | Reference |
| --- | --- |
| pGL4-hGal3p(-2638/+52) | Present work |
| pGL4-hGal3p(-1925/+52) | Present work |
| pGL4-hGal3p(-1442/+52) | Present work |
| pGL4-hGal3p(-535/+52) | Present work |
| pGL4-hGal3p(-428/+52) | Present work |
| pGL4-hGal3p(-328/+52) | Present work |
| pGL4-hGal3p(-228/+52) | Present work |
| pGL4-hGal3p(-117/+52) | Present work |
| pGL4-hGal3p(-97/+52) | Present work |
| pGL4-hGal3p(-77/+52) | Present work |
| pGL4-hGal3p(-37/+52) | Present work |
| pGL4-hGal3p(-17/+52) | Present work |
| pGEM^®^-T Vector | Promega ^®^ |
| pGL4.20 | Promega ^®^ |
| pGL4.74[hRluc/TK] | Promega ^®^ |
| pcDNA3(+)-SOX2 | Present work |
| pcDNA3(+)-SOX4 | Present work |
| pcDNA3(+)-SOX5 | Present work |
| pcDNA3(+)-SOX6 | Present work |
| pcDNA3(+)-SOX9 | Present work |
| pcDNA™3.1 (+) | Thermo Fisher Scientific |

**Supplementary Table S2.** Primers.

| Name^[2]^ | Nucleic acid target | Purpose | Sequence^[3]^ |
| --- | --- | --- | --- |
| hGAPDH-fw-RT | cDNA | Realtime PCR | TTCCACCCATGGCAAATTCC |
| hGAPDH-rv-RT | cDNA | Realtime PCR | ATCTCGCTCCTGGAAGATGG |
| RT-hSOX2-fw | cDNA | Realtime PCR | AGCAACGGCAGCTACAGCAT |
| RT-hSOX2-rv | cDNA | Realtime PCR | CTCACGTCGTAGCGGTGCAT |
| RT-hSOX4-fw | cDNA | Realtime PCR | CGACCTGCTCGACCTGAACC |
| RT-hSOX4-rv | cDNA | Realtime PCR | GCGTGCAGTAGTCCGGGAAC |
| RT-hSOX5-fw | cDNA | Realtime PCR | GCAACAGGCGGCAGGAAATG |
| RT-hSOX5-rv | cDNA | Realtime PCR | ACACCAGCAGTGGCAATGGG |
| RT-hSOX6-fw | cDNA | Realtime PCR | GGGAGTCTTGCCGATGTGGT |
| RT-hSOX6-rv | cDNA | Realtime PCR | TCTGCCAGGCTCTCAGGTGT |
| RT-hSOX9-fw | cDNA | Realtime PCR | ACCAGTACCCGCACTTGCAC |
| RT-hSOX9-rev | cDNA | Realtime PCR | GCGCCTTGAAGATGGCGTTG |
| hGal3p-for-EcoRI | Plasmid | Cloning of hGal-3p | TCAGCGAATTCTTCACTCAGGTGA |
| hGal3p_rev_HindIII | Plasmid | Cloning of hGal-3p | CTAGAAGCTTCGCTCGTTGGCTGGCTCC |
| hGal3p_for_NheI(2) | Plasmid | Long-range PCR for hGal-3p | CTAGGCTAGCTCCTTTCTCTCTGGTTCTATTTTCC |
| hGal3p-rev_EcoRImut | Plasmid | Long-range PCR for hGal-3p | GTGAAGAATTCGCTGAGAGGAATCCAAT |
| Gal3p_-1925-for_NheI | Plasmid | Deletions of hGal-3p | CTAGGCTAGCAGGGATGTGACTTTGGATTGG |
| Gal3p_-1442-for_NheI | Plasmid | Deletions of hGal-3p | CTAGGCTAGCATAAATTGGTTGCCAGGC |
| Gal3p_-535-for_NheI | Plasmid | Deletions of hGal-3p | CTAGGCTAGCAAGGTGAAGTTAAAAGGAGGG |
| Gal3p_-428-for_NheI | Plasmid | Deletions of hGal-3p | CTAGGCTAGCGAGACTAGGCCTATTGATCTA |
| Gal3p_-328-for_NheI | Plasmid | Deletions of hGal-3p | CTAGGCTAGCGCAACCTATTAATTGAATTCCTC |
| Gal3p_-228-for_NheI | Plasmid | Deletions of hGal-3p | CTAGGCTAGCACGGAACCTAACGGTGGC |
| Gal3p_-117-for_NheI | Plasmid | Deletions of hGal-3p | CTAGGCTAGCGGGCTCGCCCAGCCTGG |
| Gal3p_-97-fw_NheI | Plasmid | Deletions of hGal-3p | CTAGGCTAGCGGGGAGAGGACTGGCTG |
| Gal3p_-77-fw_NheI | Plasmid | Deletions of hGal-3p | CTAGGCTAGCAGGGGCGCCGC |
| Gal3p_-37-fw_NheI | Plasmid | Deletions of hGal-3p | CTAGGCTAGCGCTGGGAGTATTTGAGGCTCG |
| Gal3p_-17-fw_NheI | Plasmid | Deletions of hGal-3p | CTAGGCTAGCGGAGCCACCGCCC |
| hSOX2_Fwd_EcoRI | cDNA | Cloning in pcDNA3+ | CGCTCTGGAATTCATGTACAACATGATGGAGAC |
| hSOX2_Rv_XhoI | cDNA | Cloning in pcDNA3+ | CATACGCTCGAGTCACATGTGTGAGAGGGGCA |
| hSOX4_Fwd_BamHI | cDNA | Cloning in pcDNA3+ | CACTCTGGGATCCATGGTGCAGCAAACCAACAATG |
| hSOX4_Rv_EcoRI | cDNA | Cloning in pcDNA3+ | CGCACGGAATTCTCAGTAGGTGAAAACCAGGTTG |
| hSOX5_Eco_for | cDNA | Cloning of human SOX5 | CTAGGAATTCATGCTTACTGACCCTGATTTACC |
| hSOX5_Xho_rev | cDNA | Cloning of human SOX5 | CTAGCTCGAGTCAGTTGGCTTGTCCTGC |
| hSOX6_EcoR_for | cDNA | Cloning of human SOX6 | CTAGGAATTCATGTCTTCCAAGCAAGCCAC |
| hSOX6_Xho_rev | cDNA | Cloning of human SOX6 | CTAGCTCGAGTCAGTTGGCACTGACAGC |
| hSOX9_EcoR_for2 | cDNA | Cloning of human SOX9 | CTACGAATTCATGAATCTCCTGGACCCCTTCATGAAG |
| hSOX9_Xho_rev2 | cDNA | Cloning of human SOX9 | CTACCTCGAGTCAAGGTCGAGTGAGCTGTGTGTAG |
| hSOX9_EcoR_for_dATG | Plasmid | Cloning of HALO-Fusion | CTACGAATTCAATCTCCTGGACCCCTTCATGAAGATG |
| hSOX9_Xba_rv | Plasmid | Cloning of HALO-Fusion | CTACTCTAGATCAAGGTCGAGTGAGCTGTGTGTA |

**Supplementary Table S3.** Real time PCR primer efficiencies.

| Primer pair | Efficiency (%) | R^2^ values |
| --- | --- | --- |
| SOX2 | 110.24 | 0.9894 |
| SOX4 | 109.62 | 0.9818 |
| SOX5 | 111.77 | 0.9933 |
| SOX6 | 108.34 | 0.9819 |
| SOX9 | 106.22 | 0.9904 |
| GAPDH | 110.83 | 0.9953 |

**Supplementary Table S5.** Top 20 potential binding sites of SOX proteins in the human *LGALS3* promoter

| Matrix ID | Binding site relative to TSS | Binding site relative to TSS | Predicted sequence | Relative score |
| --- | --- | --- | --- | --- |
| MA0442.3.SOX10 | -1698 | -1693 | ACAAAG | 1.00 |
| MA0442.1.SOX10 | -1698 | -1693 | CTTTGT | 1.00 |
| MA1152.2.SOX15 | -1833 | -1827 | CTTTTGT | 0.98 |
| MA1152.2.SOX15 | -2243 | -2237 | TTTTTGT | 0.95 |
| MA1152.2.SOX15 | -2050 | -2044 | TTTTTGT | 0.95 |
| MA1152.2.SOX15 | -545 | -539 | TTTTTGT | 0.95 |
| MA0867.3.SOX4 | -1833 | -1826 | AACAAAAG | 0.95 |
| MA1563.2.SOX18 | -300 | -293 | AAGAATC | 0.95 |
| MA0084.2.SRY | -2048 | -2042 | AAACAAA | 0.95 |
| MA0867.2.SOX4 | -2244 | -2235 | GAACAAAAAG | 0.94 |
| MA1152.1.SOX15 | -2243 | -2234 | TTTTTGTTCT | 0.94 |
| MA0867.3.SOX4 | -2243 | -2236 | AACAAAAA | 0.94 |
| MA0867.3.SOX4 | -2050 | -2043 | AACAAAAA | 0.94 |
| MA0867.3.SOX4 | -546 | -539 | AACAAAAA | 0.94 |
| MA0867.2.SOX4 | -1834 | -1825 | TAACAAAAGG | 0.93 |
| MA0084.2.SRY | -1831 | -1825 | TAACAAA | 0.93 |
| MA0084.2.SRY | -917 | -911 | TAACAAA | 0.93 |
| MA1152.2.SOX15 | -349 | -343 | GTTTTGT | 0.93 |
| MA1152.1.SOX15 | -1833 | -1824 | CTTTTGTTAG | 0.93 |
| MA0084.2.SRY | -1743 | -1737 | ATACAAT | 0.93 |

**Supplementary Table S6.** Top 20 TBFS in the -117/+52 region of the human *LGALS3* promoter

| Matrix ID | Binding site relative to TSS | Binding site relative to TSS | Predicted sequence | Relative score |
| --- | --- | --- | --- | --- |
| MA1513.2.KLF15 | -44 | -37 | CCCCGCCC | 1.00 |
| MA0493.3.KLF1 | -44 | -37 | GGGCGGGG | 1.00 |
| MA1959.2.KLF7 | -44 | -37 | GGGCGGGG | 1.00 |
| MA2341.1.FEZF2 | -110 | -103 | CCCAGCCT | 1.00 |
| MA1513.1.KLF15 | -46 | -36 | GCCCCGCCCGG | 0.97 |
| UN0628.1.ZNF543 | -75 | -64 | GGGGCGGCGCCC | 0.96 |
| MA0516.3.SP2 | -67 | -59 | GAGGCGGGG | 0.96 |
| MA1961.1.PATZ1 | -47 | -36 | GCCGGGCGGGGC | 0.96 |
| MA1961.2.PATZ1 | -46 | -36 | CCGGGCGGGGC | 0.96 |
| MA1961.2.PATZ1 | -68 | -58 | CGAGGCGGGGC | 0.95 |
| MA1961.1.PATZ1 | -68 | -57 | CCGAGGCGGGGC | 0.95 |
| MA2328.1.ZBED4 | -71 | -62 | GCCGCCCCGC | 0.95 |
| MA0516.1.SP2 | -59 | -45 | GGCCCGCCTCTCCCG | 0.94 |
| UN0635.1.ZNF571 | -16 | -2 | GCGGGGCGGTGGCTC | 0.93 |
| UN0635.2.ZNF571 | -16 | -4 | GGGGCGGTGGCTC | 0.92 |
| UN0593.2.ZNF100 | -14 | -2 | GCGGGGCGGTGGC | 0.91 |
| UN0635.1.ZNF571 | -76 | -62 | GCGGGGCGGCGCCCC | 0.91 |
| UN0593.1.ZNF100 | -16 | -2 | GCGGGGCGGTGGCTC | 0.89 |
| MA1713.1.ZNF610 | -74 | -61 | GGCGCCGCCCCGCC | 0.89 |
| UN0666.1.ZNF891 | -22 | -5 | GGGCGGTGGCTCCGAGCC | 0.87 |

1. Concentrations were measured by Implen 300 photometer and purity checked, making sure the 260/280 value is around 1.8.
2. Name includes in all cases the direction of the primer (fw: forward, rv: reverse) and gene of interest.
3. Primers were designed manually and ordered at Metabion.
